# Genetic heterogeneity in *Taenia solium* cysticerci among naturally infected pigs revealed by microsatellite markers and associated epidemiological risk factors

**DOI:** 10.1101/2025.05.21.655296

**Authors:** Rashmi Sharma, Rabinder Singh Aulakh, Balbir Singh Bagicha, Rajnish Sharma, Randhir Singh

## Abstract

**Background:** The tapeworm, *Taenia solium*, also called as pork tapeworm is acquired by human on consumption of raw or undercooked pork infested with intermediate larval stage. The parasite is known to establish an obligatory cyclozoonoses with human serving as the only definitive host for the organism. India is endemic for the presence of *T. solium* infection particularly for cysticercosis both in human and pig population.

**Material method:** A total of 389 pigs were screened from Ludhiana, Amritsar, Bathinda, Jalandhar, Patiala and in union territory, Chandigarh were screened for *T*. solium. The collected samples were analyzed in relation to different epidemiological variables – Location, age, sex, breed, and management system and to conduct a PCR targeting valine t-RNA and NADH subunit-2. Against molecular confirmed *T. solium*, genetic polymorphism was asses for seven microsatellite satellite markers.

**Findings:** A prevalence rate 5.14% on post mortem examination as well as molecular confirmation of suspected samples (PCR-targeting NADH subunit-2 gene). Highest prevalence rate was observed in Patiala (15%) with lowest in Bathinda (0.00%), due to low participation of the pig farmers of that region. We also observed genetic variability for seven microsatellite markers with TSSR_09 (26/200) and TSSR_01 (14/200) were the 2 microsatellites showing highest variability.

**Conclusion:** *T. solium* cysticercosis is a neglected tropical disease with significant public health hazard. Measures should be taken to prevent transmission of pathogen as well as sale of infected pork.

## Introduction

*Taenia solium* is a major public health problem, associated with pork consumption in developing countries (1) and is potentially an emerging parasitic disease in Europe (2). It is a leading cause of death from foodborne infections, as per WHO Foodborne Disease Burden Epidemiology Reference Group, 2015 (3). Infection with Cysticercus cellulosae (larval stage) is manifested by ingestion of worm eggs. The situation is grave since thousands of them are produced at a time and thus can establish neurocysticercosis in humans in more numerous numbers, which mostly ends critically (4). There are three known zoonotic species of the genus *Taenia*-*T. solium, T. saginata* and *T. Asiatica*, however *T. solium* is the most significant zoonotic species among them.

World Health Organization (WHO) along with Food and Agricultural Organization (FAO) has declared *T. solium* as a parasite of “greatest global concern” (5). The disease is endemic in south and south-east asia and is an emerging problem in certain parts of sub-Saharan Africa; it is a serious problem for decades in and around Latin America (6).

*Taenia solium* infection has two clinical forms in humans-taeniasis and cysticercosis. *T. solium* taeniasis in humans develops after 6-8 weeks post-ingestion of cysticerci and is characterized by non-specific mild signs like abdominal pain, nausea, diarrhoea and constipation, even in case of heavy infection which resolve on its own on (7;8).

Cysticercosis has a variable incubation period and people can be asymptomatic for years. The disease develops on accidental intake of food and/ or water contaminated with eggs of *Taenia spp*. and depending upon the location of cyst development. It can be categorized as generalized infection (cyst present throughout the body), subcutaneous cysticercosis (cyst localized in the subcutaneous tissue), ophthalmo-cysticercosis and neurocysticercosis (affecting eyes, cranial and sub-cranial region respectively) (9).

Neurocysticercosis is the most avertable and the leading cause of epilepsy throughout the world. It is responsible for 30% of epilepsy cases in endemic areas (6). Other symptoms observed during the disease are chronic headaches, blindness, seizures, meningitis, hydrocephalus, focal neurological deficits, psychological disorders, dementia, ocular and spinal cysts (10; 11, 12;13).

Cysticercosis in pigs shows a dissimilar pattern of being asymptomatic in nearly all the observed cases, which could be due to their limited lifespan because of early slaughter, however, pigs with neurocysticercosis have been reported to develop clinical signs. These could be autonomic in nature (like chewing motions along with increased salivation and ear stiffening), motor signs such as tonic seizures even tremors and stereotypical circle walking (14).

Familiarity or data regarding the genetic build of *T. solium* can serve as the very basis of molecular epidemiology for the parasite. It can be helpful in determining the transmission of the diseases and variation in the infectivity potential between strains (13). The use of microsatellite markers will be helpful in determining their heterozygosity at the local community level. Genetic tools facilitate better understanding of parasitic interaction with porcine and human transmission.

Additionally, it may be resourceful in determining the route of transmission in case of an outbreak and pinpointing the source of infection or the area of origin. There’s scarcity of data in relevance to the genetic diversity of organisms in the country, also studies performed in *T. solium* cysticerci recovered from pigs with different geographical origins have shown that genetic variations of the parasite may be involved in pathogenicity. These finding prompted us to devise the epidemiological study to assess the genetic variation of *T. solium* in naturally infected pigs.

## Material Method

The objective of this experiment was to determine molecular prevalence of *T. solium cysticercosis*, asses genetic variability in naturally infected pigs and the association of epidemiological factors in its occurrence.

### Ethics Statement

The present study was ethically approved by Institutional Ethics Committee of Guru Angad Dev Veterinary and Animal Sciences University, Ludhiana from January, 2020 till December, 2021 as per IAEC/2020/81-110.

### Study Site

Punjab is in north western region of India, has an agrarian economy. As per 20^th^ livestock census, Punjab has 52000 pigs, with pig farming an emerging industry. Slaughter practices in Punjab are largely informal, with the only official slaughterhouse located in the Union Territory of Chandigarh. For the study, pigs were screened from the randomly selected five district of the state-Ludhiana, Amritsar, Bathinda, Jalandhar, Patiala and in union territory, Chandigarh.

### Target population

Pig population reared in intensive, semi-intensive and extensive farming system in 5 districts of Punjab was target the population. Pigs found positive for T. solium cysticercosis.

### Sample size

A sample size of 385 was estimated using statulator (15) at 95% level of confidence, and 5% margin of error for sample size and assuming an expected rate of prevalence at 5%.

### Sample collection

An initial visit was conducted to the local meat slaughtering shops and two shops were selected from each district for even representation of different districts. Subsequently, regular visits were made to the selected meat shops where slaughtered pigs were screened for *T. solium* cysticercosis. In addition, 55 pigs were sampled in 3 visits from the official slaughter house in state capital. The distribution of samples collected was analyzed in relation to different epidemiological variables – Location, age, sex, breed, and management system, as provided below in table 1. The occurrence level of infection with respect to selected risk factors was calculated in sampled pig population.

**Table 1:** Details of the samples collected as per different epidemiological variables.

| **Risk Factors** |  |  |
| --- | --- | --- |
| **S. No.** | **District (Tehsil)** | **No. of Sample screened** |
| 1 | Amritsar (Ajnala) | 109 |
| 2 | Bathinda (Bathinda) | 44 |
| 3 | Jalandhar (Jalandhar) | 90 |
| 4 | Ludhiana (Ludhiana West) | 70 |
| 5 | Patiala (Rajpura) | 20 |
| 6 | Chandigarh (Chandigarh) | 55 |
| **Gender** |  |  |
| 1 | Male | 108 |
| 2 | Female | 281 |
| **Breed** |  |  |
| 1 | Desi | 118 |
| 2 | White Yorkshire | 278 |
| **Management System** |  |  |
| 1 | Free-range | 87 |
| 2 | Semi-intensive | 198 |
| 3 | Intensive | 104 |

### Necropsy

On post examination, full carcass dissections were performed, tongue, masseter muscle, thigh muscles, diaphragm, heart muscles and brain were visually examined. Part of positive pigs’ carcasses were collected in sterile biohazard bags and transported to the lab in a maintained cold chain. An estimation of number of viable cysts from 250gm of infected muscle tissue was done.

### Collection of cysts for microscopic examination

The collected samples were first analyzed microscopically in the laboratory. Positive pork samples were incised carefully, fiber by fiber and viable cysts were separated and placed in petridish with water. A single cyst from the petridish was taken between the two clean glass slides, gently pressed and then visualized under the microscope for the presence of hooks and suckers.

### Collection of cysts for DNA extraction

Ten healthy and viable cysts were randomly collected from each positive pork samples. The cysts were washed in normal saline, to remove any pork residue. The cysts were individually homogenized using tissue homogenizer (Tissue Lyser II, Qiagen). DNA extraction was carried out using DNeasy blood and tissue kit as per manufacturer’s guidelines. The DNA quality and quantity was determined by using UV-spectrophotometry (Nanodrop-2000, ThermoScientific) and eluted DNA was stored at -20°C until further use.

#### 3.3.9 Multiplex Polymerase Chain Reaction (PCR)

The oligonucleotide sequence used for multiplex PCR amplification targeted *T. solium* valine t-RNA and NADH (subunit-2) published in literature (16) and listed in table 2. The selected oligonucleotides enabled species differentiation and confirmation between *T. solium, T. saginata and T. asiatica*. All primers for the assay were obtained from EUROFINS Ltd.

**Table 2:**
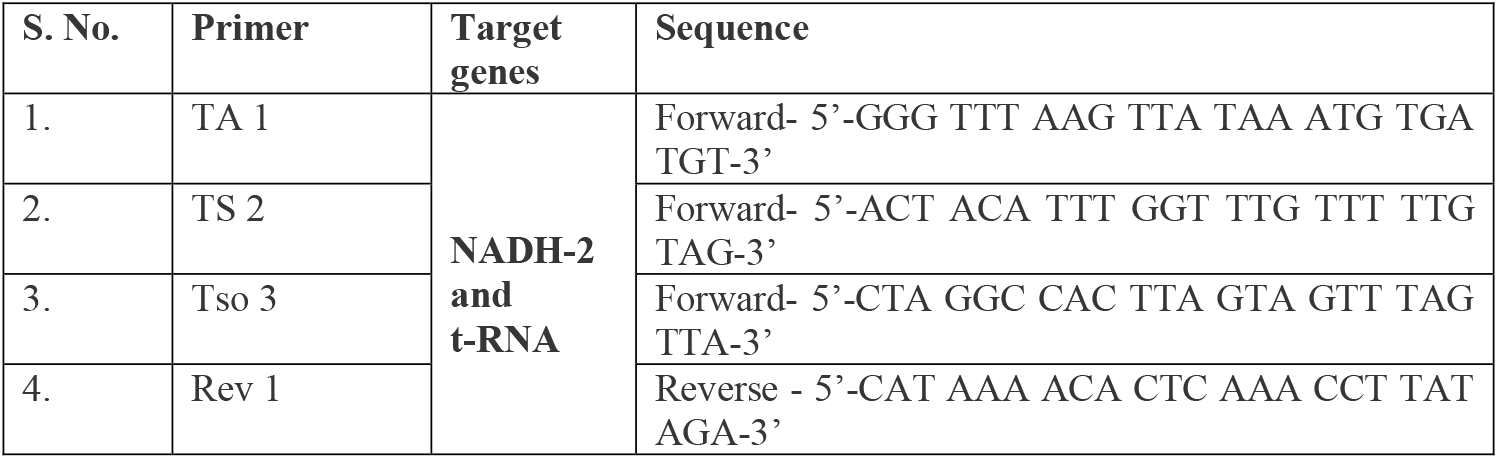
Detailed information of primer sequences used in multiplex PCR.

The PCR reaction was carried out with a total reaction volume of 25μl consisting 12.5μl of master mix (GoPro master mix), 1μl each of forward and reverse primer, 4μl of cyst extracted DNA and 7.5μl of nuclease free water (NFW). The thermal cyclic condition used for speciation of *Taenia* spp. included initial denaturation of 5 minutes at 94 ºC followed by 35 amplification cycles (denaturation at 94 **º**C for 30 seconds, annealing at 54 **º**C, extension at 72 **º**C) and final extension at 72 **º**C for 7 minutes in Master cycler gradient Thermocycler’s (Eppendorf and AB system). Morphological confirmed *T. solium* DNA (microscopically verified) were used as positive control whereas NFW was used as negative control.

### Microsatellite analysis

After initial species confirmation of extracted *T. solium* metacestodes, microsatellite PCR was performed targeting Seven microsatellite loci-TSSR_01, TSSR_09, TSSR_16, TSSR_18, TSSR_27, TSSR_28 and TSSR_32. The oligonucleotide sequences were taken from previously published data (17; table 3). All the reactions were carried out using Master cycler gradient Thermocycler’s (Eppendorf and AB system). Primer sequences were procured from Eurofins (Eurofins Scientific, Europe) and listed in table 3. The PCR reaction volume of 25μl included 12.5μl master mix, 1μl each of forward and reverse primer, 4μl of cyst extracted DNA and 7.5μl of NFW. Modifications were made for TSSR 27, 0.5ml of MgCl_2_ and nuclease free water was adjusted to 7μl. For TSSR_28, 12.5μl Type -it PCR master mix solution was used with 2.5 μl each of forward and reverse primer, 5 μl of Q solution, template DNA 3μl and 1.5 μl of RNase free water.

**Table 3:**
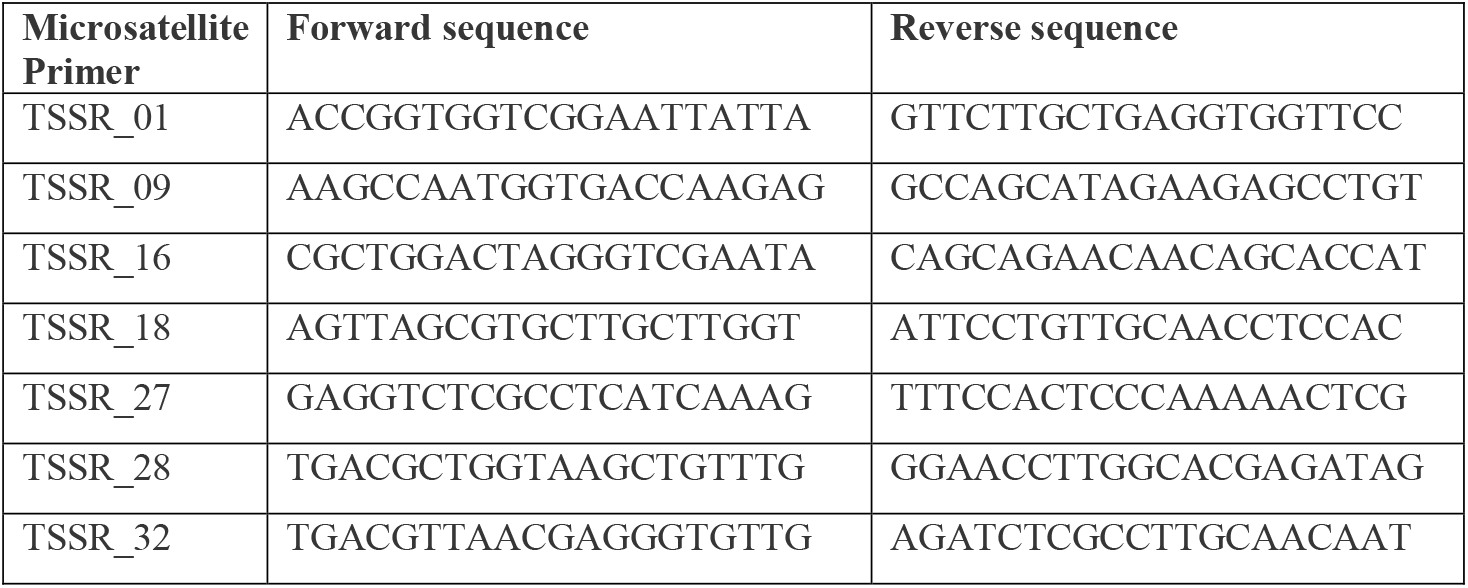
Primer sequences use for Microsatellite PCR.

The thermal cyclic conditions for TSSR 01, TSSR 09, TSSR 16, TSSR 18 and TSSR 32 consisted of an initial denaturation of 94 ºC for 5 minutes, 30 cycles of amplification (cyclic denaturation at 94 ºC for 30sec, annealing at a gradient of 52-56 ºC for 30 sec and elongation at 72 ºC for 30sec) followed by final elongation at 72ºC for 7 minutes. For TSSR_27, protocol was modified to include 35 cycles and the annealing step was extended to 45 sec. The thermal cyclic conditions for TSSR_28 included initial denaturation is done at 95 ºC for 5 minutes followed by 28 cycles (cyclic denaturation at 95 ºC for 30sec, annealing at 58 ºC and elongation at 72 ºC for 45 sec) and concluded with a final elongation at 60 ºC for 30 minutes.

The initial speciation PCR product (474 bp) and PCR products obtained for microsatellite markers were visualized using low melting point agarose gel (Qiagen, Germany) at a concentration of 1.5% for the speciation product and 2% for the purpose of gel electrophoresis (Tarson, Kolkata). Five microliter of product was loaded into the gel wells and 1X TBE buffer was used and DNA bands were visualized using a gel documentation system.

### Statistical analysis

The apparent and true prevalences were estimate using Epi Tools with estimated 95% confidence interval. Chi-square test was done to assess the effect of epidemiological variables on disease prevalence with the help of Ms-excel, 2019.

### 4.3 Prevalence of *Taenia solium* cysticercosis in pigs of Punjab

Out of 389 pig carcasses that were examined, 20 were found positive indicating an overall prevalence of 5.14%. Overall, highest apparent prevalence was recorded in Patiala with 15% (5.24%-36.04%) followed by Jalandhar (7.78%, 3.82%-15.20%), Ludhiana (7.17%, 3.09%-15.65%) and least number of carcasses were positive in Amritsar (2.25%, 0.94%-7.78%) on post mortem examination. None sample was positive from Bathinda district. The study confirmed the presence of porcine cysticercosis in the state of Punjab with relatively higher prevalence in unorganized slaughter shops of Punjab.

In present study, 7.40% (8/108, as shown in table in 28) males and 4.27% (12/28, as shown in Table 29) females were found positive and prevalence rate recorded were 5.93% in desi breed of pigs whereas for white Yorkshire 4.27% animals were positive. Desi breed of pigs had higher rate of infection (5.93%) with semi-intensive system of rearing responsible for maximum exposure to *T. solium* infection. No statistical significance was observed amongst disease occurrence and various epidemiological factors (p>0.05). Detailed information on district wise prevalence rate is provided in table 4 (95% CI).

**Table 4:** Apparent area wise prevalence of *Taenia solium cysticercosis* in pigs of Punjab.

| S. No. | Sampling Area | No. of samples collected | No. of positive sample | Apparent prevalence | 5% Confidence interval (CI) | True prevalence | 5% Confidence interval (CI) |
| --- | --- | --- | --- | --- | --- | --- | --- |
| **Area** |  |  |  |  |  |  |  |
| 1 | Amritsar | 109 | 3 | 2.25% | 0.94%-7.78% | 1.97% | 0.56%-6.65% |
| 2 | Bathinda | 44 | 0 | 0.00% | 0.00% | 0.00% | 0.00% |
| 3 | Jalandhar | 90 | 7 | 7.78% | 3.82%-15.20% | 7.62% | 3.71%-15.00% |
| 4 | Ludhiana | 70 | 5 | 7.14% | 3.09%-15.65% | 6.90% | 2.94%-15.34% |
| 5 | Patiala | 20 | 3 | 15.00% | 5.24%-36.04% | 15.73% | 5.63%-38.88% |
| 6 | Chandigarh | 55 | 2 | 3.64% | 1.00%-12.33% | 2.97% | 0.73%-11.35% |
| **Sex** |  |  |  |  |  |  |  |
| 1 | Male | 108 | 8 | 7.40% | 3.80%-13.94% | 7.20% | 3.66%-13.68% |
| 2 | Female | 281 | 12 | 4.27% | 2.46% - 7.31% | 3.67% | 2.02%- 6.57% |
| **Breed** |  |  |  |  |  |  |  |
| 1 | Desi | 118 | 7 | 5.93% | 2.90%-11.74% | 5.54% | 2.64%-11.24% |
| 2 | White Yorkshire | 278 | 13 | 4.63% | 2.73%- 7.76% | 4.08% | 2.32%- 7.08% |
| **Management system** |  |  |  |  |  |  |  |
| 1 | Free-range | 87 | 4 | 4.60% | 1.80%-11.24% | 4.04% | 1.49%-10.47% |
| 2 | Semi-intensive | 198 | 11 | 5.56% | 3.13%-9.68% | 5.12% | 2.82%-9.13% |
| 3 | Intensive | 104 | 5 | 3.85% | 1.51%- 9.48% | 3.20% | 1.15%-8.58% |

The morphological analysis of *Cysticercus cellulosae* revealed morphological variation in appearance of cyst found in different areas of carcass. Cysts in tongue were large and opaque with reddish tinge (fig.1A) whereas those observed in muscles were significantly smaller (fig. 1B). Neurocysticercosis was observed in pigs, the cyst was small and has an everted scolex unlike inverted in others (fig.1C). The cyst extracted had a translucent outer wall, inside which a scolex and cystic fluid (fig.1B). on microscopic examination of cyst, scolex with four suckers and double rows of rostellar hook pattern was seen and hooks were anvil shaped with handle, guard and hook. (fig. 1D & 1E).

**Fig. 1A.**
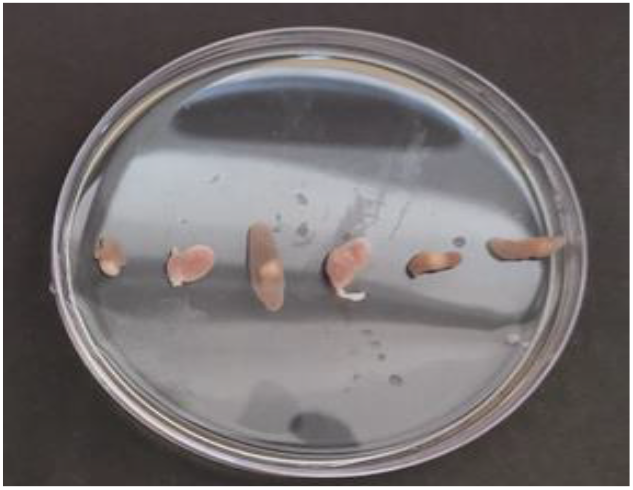
*C. cellulosae* in tongue

**Fig. 1B.**
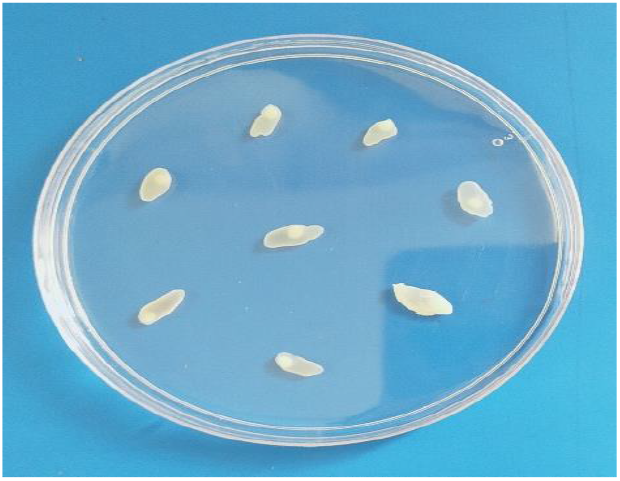
*C. cellulosae* in muscle

**Fig. 1C.**
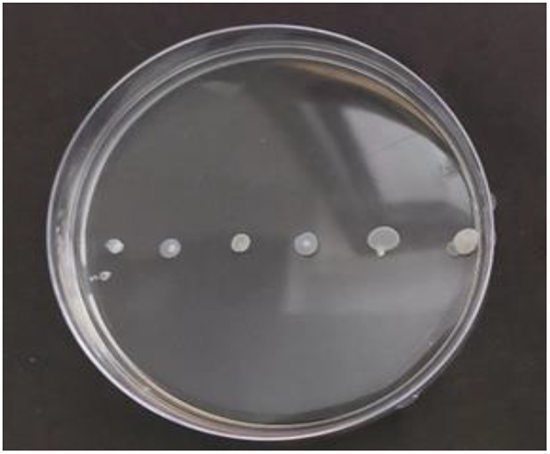
*C. cellulosae* from brain

**Fig. 1D.**
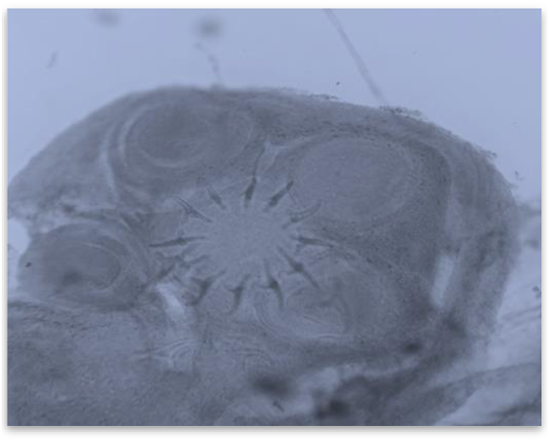
Scolex with suckers and hook

**Fig. 1E.**
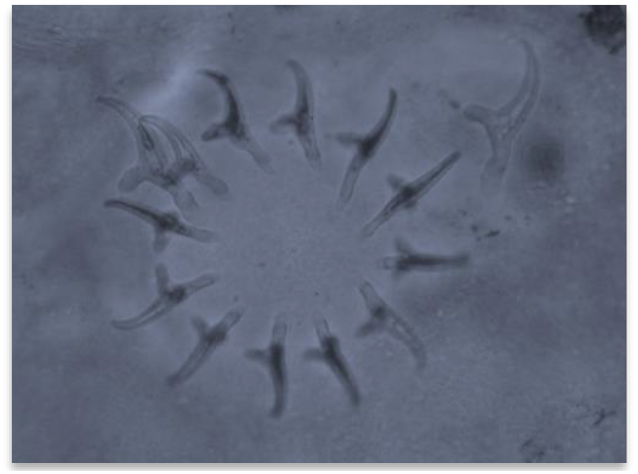
Anvil shaped hook of *C. cellulosae*

### Molecular characterization of *Taenia solium* cysticercosis

From each of the positive sample’s DNA was extracted from 10 cyst samples and all these 200 DNA samples were then screened by using multiplex PCR reaction targeting NADH subunit-2 and t-valine genes. Molecular prevalence overall as well as in terms of area-related was same as that of prevalence observed through carcass examination. A product size of 474bp observed for all 200 cysts as shown figure 2.

**Figure 2:**
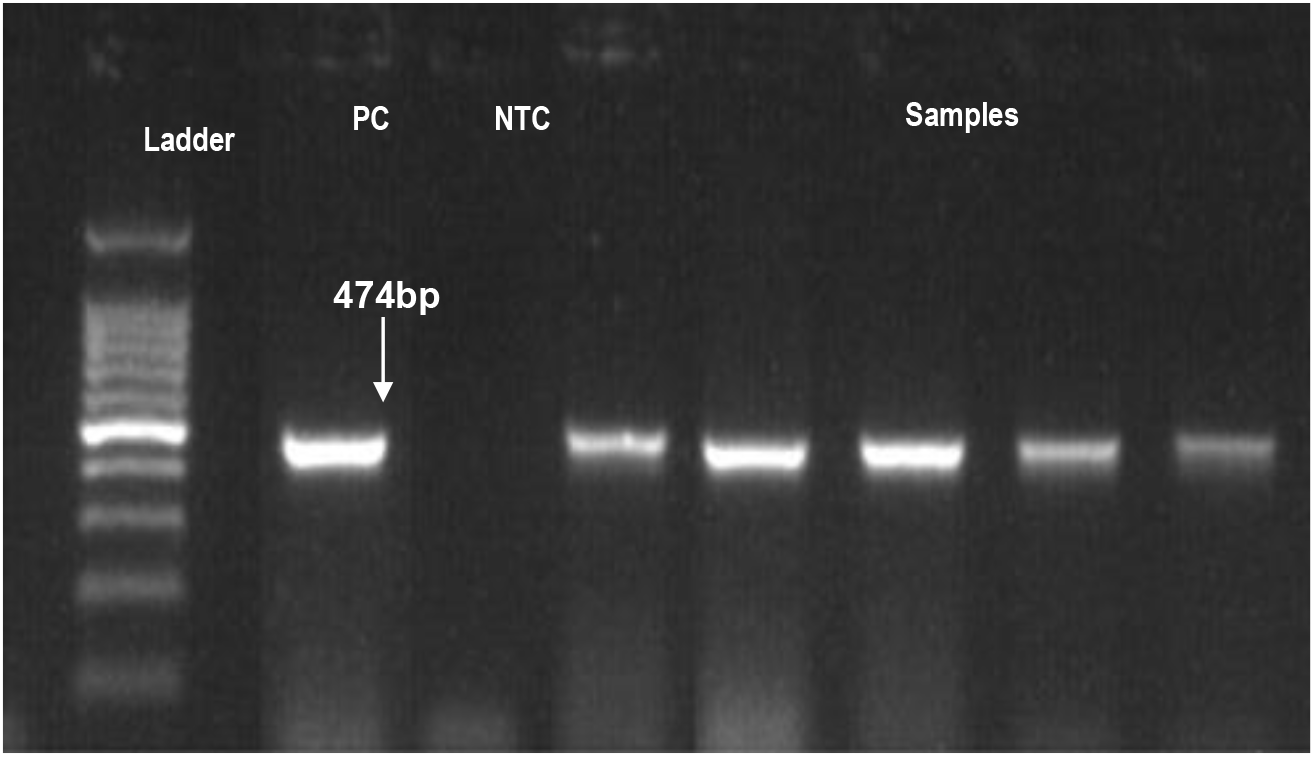
Gel electrophoresis of products of multiplex PCR from pork samples Sequencing and phylogenetic analysis.

The phylogenetic tree alignment based on NADH subunit-2 depicts the isolates are clustered around *T. solium* suggesting that sample ID 45, 101, 191, 202, 1710, 1810 are the strains of the same species as shown in figure 3.

**Fig 3.**
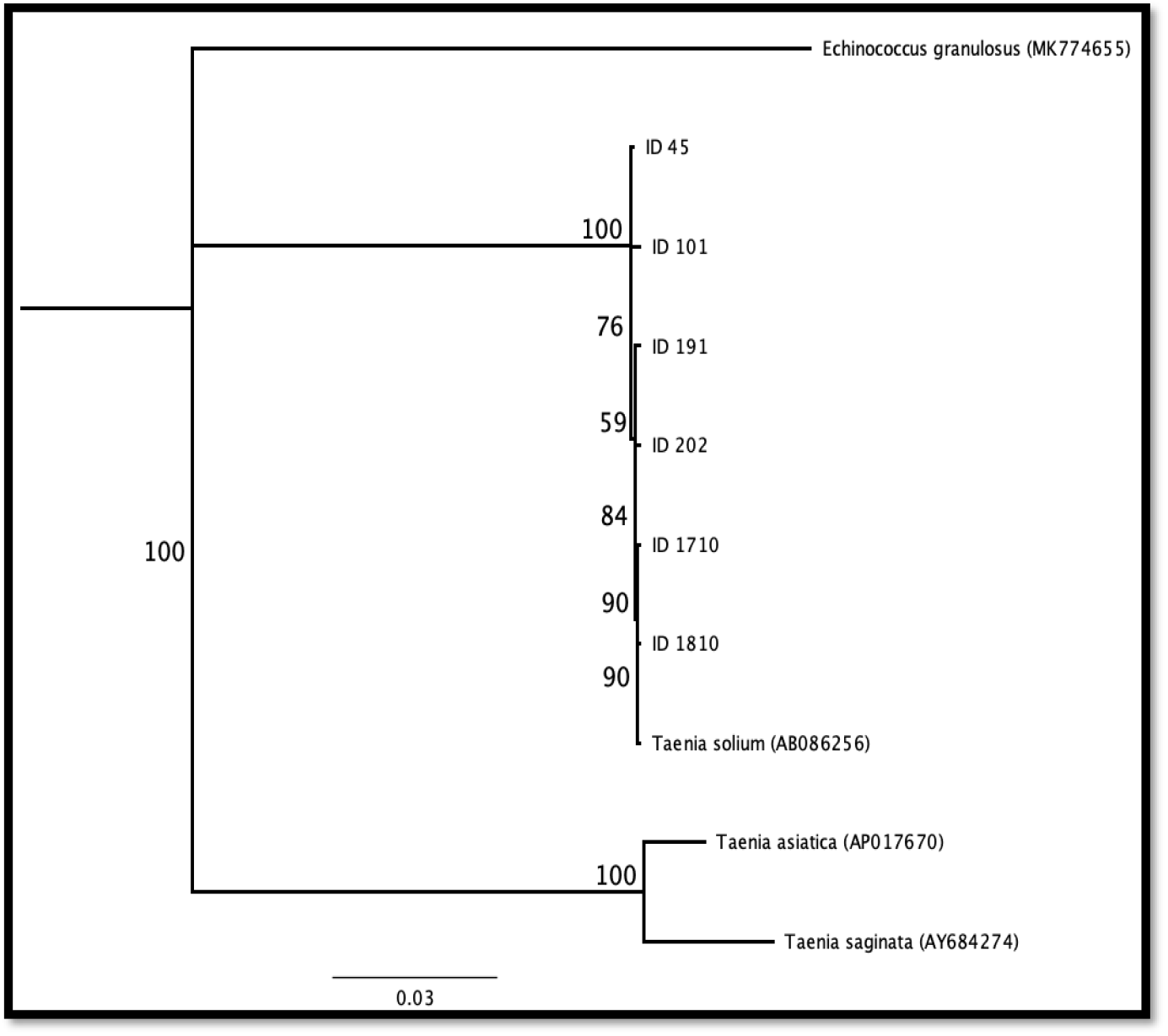
Phylogenetic Tree of t-RNA and NADH-2 of *Taenia solium* isolates from naturally infected pigs and reference strains.

### Microsatellite analysis of *Taenia solium* metacestode

All the confirmed 200 C. *cellulosae* DNA was used to assess the genotypic variations using short tandem repeats (STRs). Seven microsatellites, TSSR_01, TSSR_09, TSSR_16, TSSR_18, TSSR_27, TSSR_28and TSSR_32 product size obtained is shown in table 5.

**Table 5:** Product size of targeted microsatellites.

| S. No. | Microsatellite Targeted | PCR product size |
| --- | --- | --- |
| 1 | TSSR_01 | ~220 |
| 2 | TSSR_09 | ~140 |
| 3 | TSSR_16 | ~150 |
| 4 | TSSR_18 | ~160 |
| 5 | TSSR_27 | ~150 |
| 6 | TSSR_28 | ~190 |
| 7 | TSSR_32 | ~180 |

Variations in certain *T. solium* cysts were observed in regards to the band pattern that was observed in majority of cysts. For TSSR_01 and TSSR_18 variants had the band size was of slightly larger size than observed normally (>220bp, and >160bp,) whereas band variation observed in TSSR_09, TSSR_16 and TSSR_28 microsatellite had a smaller than normal shown size (<140bp, Plate 17, <150bp, Plate 18 and <190bp. In microsatellites TSSR_27 and TSSR_32, double band pattern was observed of band size of 150bp and 180bp (fig. 4).

**Fig 4.**
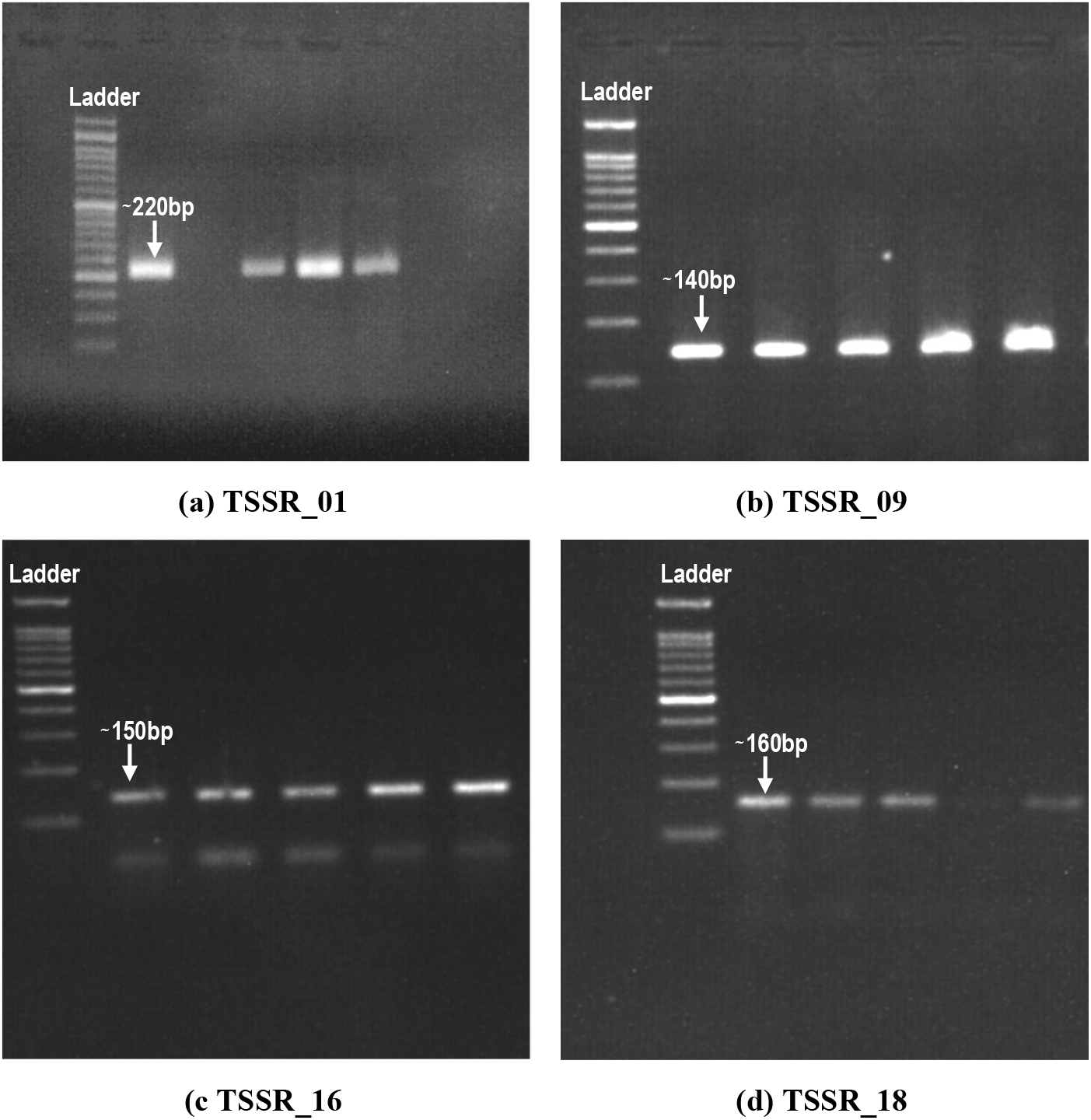
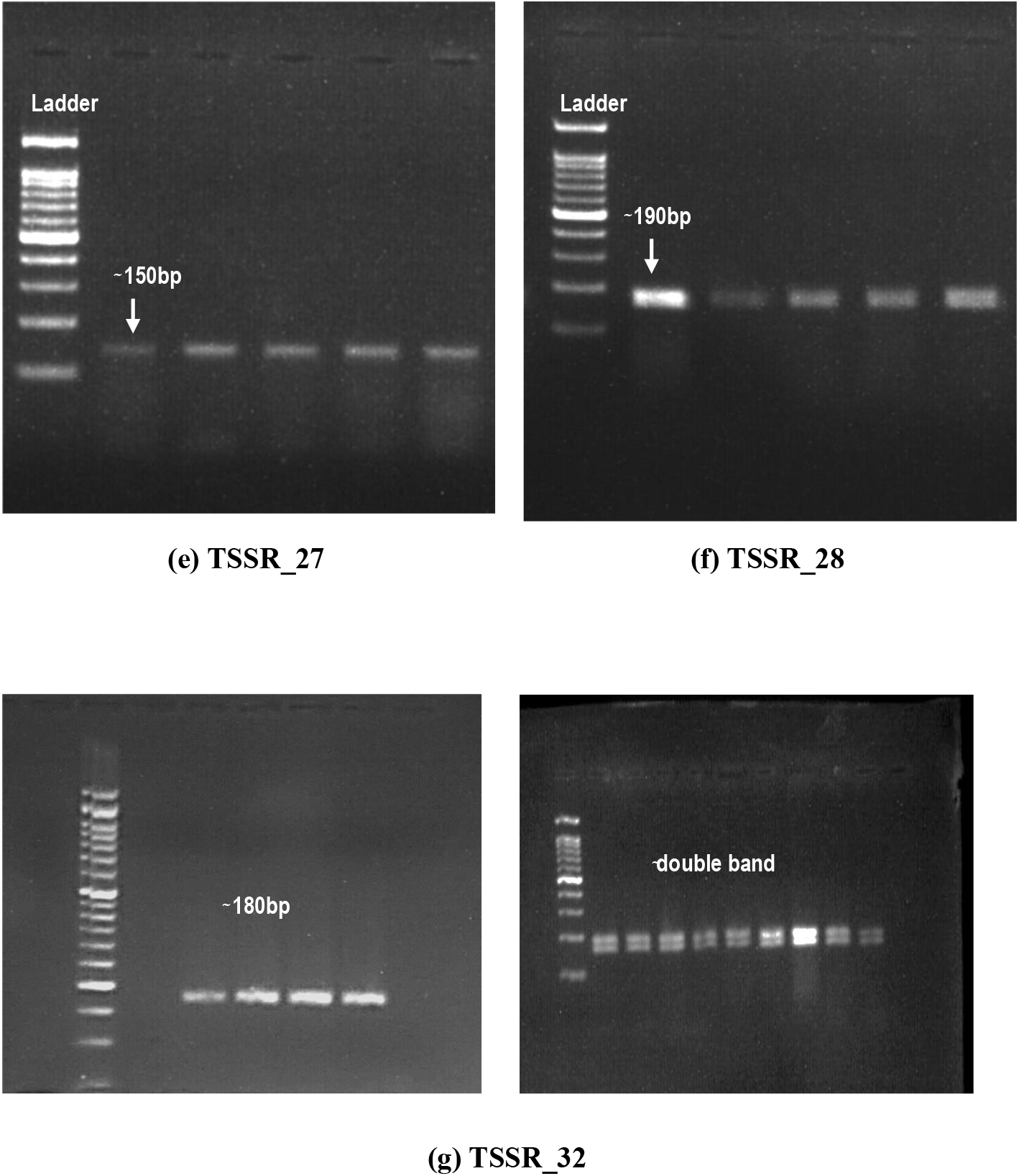
Gel Electrophoresis visualization of microsatellite product.

Variation rate observed in Amritsar and Jalandhar was equal (63.33%, 19/30 and 39/70 for Amritsar and Jalandhar respectively), Ludhiana (57.50%, 23/40), Patiala (46.67%, 14/30) and Chandigarh (70.00%, 14/20) as shown in figure 5. A total 56% (112/200) cysts demonstrated variations with 50% (100/200) cysts showing one band variation and 6% (12/200) variation were shown in double bands.

**Fig 5.**
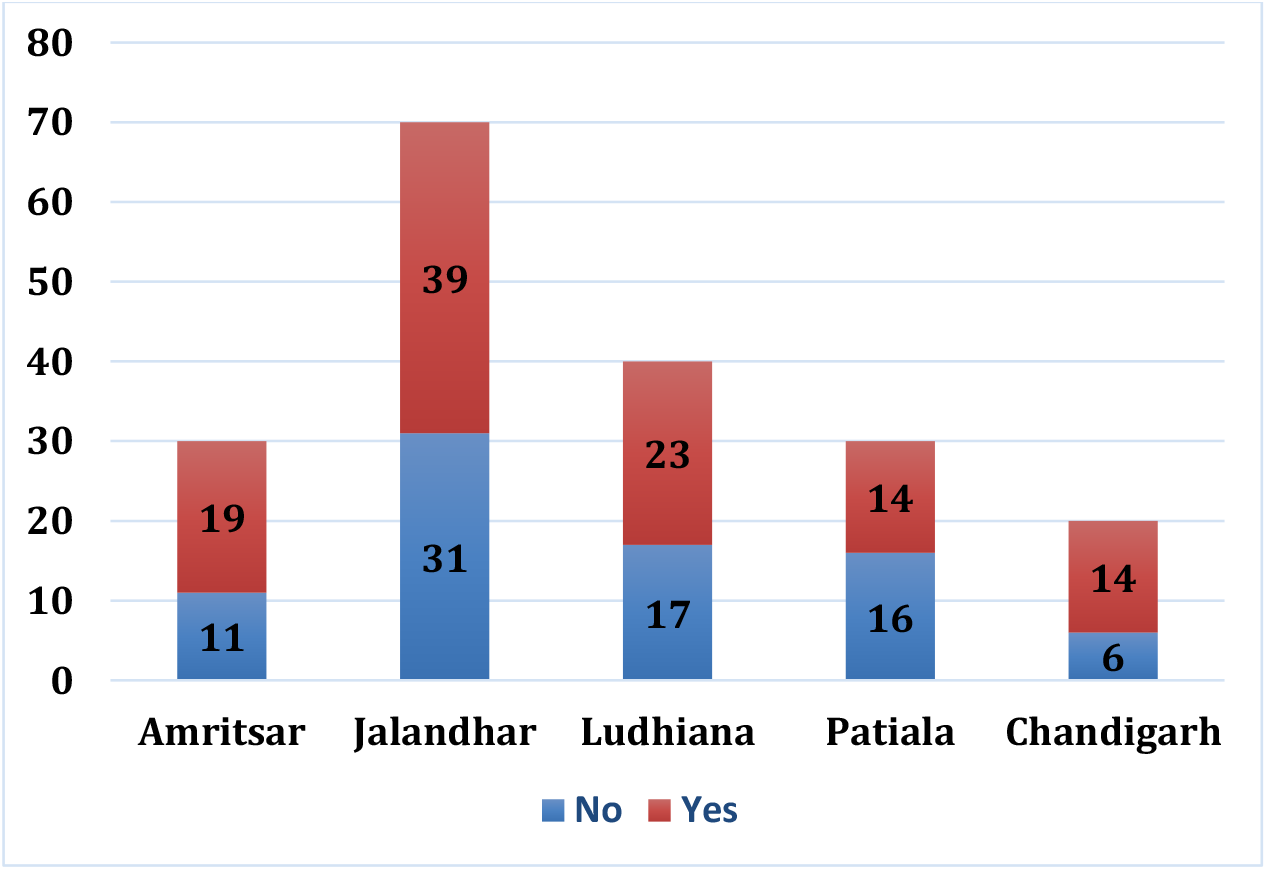
Band Variation observed in Districts.

Highest number of variation was observed in TSSR_32 (60/200) followed by TSSR_09 (26/200) followed by TSSR_01 (14/200) and least was seen in TSSR_16 (3/200) as shown in figure 6.

**Fig. 6:**
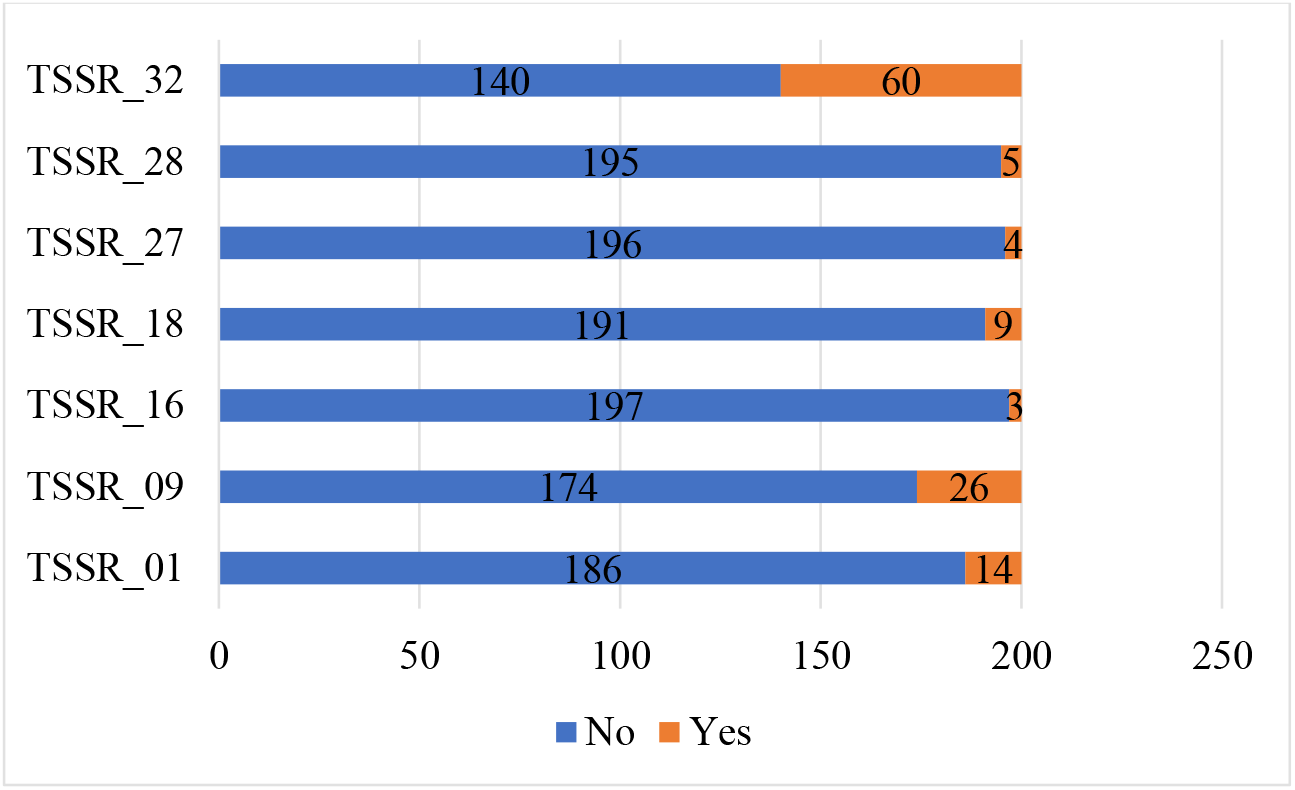
Band Variation in microsatellites.

## Discussion

The present study confirms the presence of *T. solium* infection in Punjab (5.14%). A similar study which estimated the prevalence rate of porcine cysticercosis in Punjab was conducted in Punjab (Singh *et al*., 2018), observed an apparent overall prevalence rate 1.40% and district wise prevalence rates were between 3.50%,3.30% and 1.18% for Jalandhar, Ludhiana and Patiala respectively, which was considerably lower than what was observed in this current study. The variation in prevalence could be because of availability as well as accessibility of the infected animal to the researcher was affected due lack of participation by the pig slaughtering shop owner. Also, seasonal variability and management practices followed for rearing purposes of the pigs could be the possible reason behind such prevalence variation. Prevalence recorded in current study is underrepresentation of data since only 5 randomly selected districts were screened and positive cases from other districts of Punjab were not screened for the study.

The current study used traditional post mortem method for detection of infection in pigs, although PM method is effective for heavily infected pigs but light infection could have been missed. Serological method of cysticercosis detection in pigs is far more accurate in detecting infected pigs regardless of severity of infection.

The current observed higher number of males to be positive (7.40%) for cysticercosis than females. A study done in Jammu (18), observed similar trends in prevalence in reference to sex as well as breed were seen, higher males (1.36%) were found to be positive than females (0.85%). Saravanan *et al*. (2014) on the other hand showed sex wise prevalence to be more in female (5.43%) than in males (4.82%). Similarly, study in Assam (20) reported prevalence of *Cysticercus cellulosae* infection in male and female was 9.15 and 10.39 per cent respectively.

Since pigs produce large litters and female can be bred as early as 6-8 months of age with an average weight of 80-100kg. Females are usually kept for breeding purpose whereas males are usually slaughter early and in more numbers. Since more males are slaughtered a skewed infection rate in them is possible. Males since considered expendable since not used for breeding are more commonly reared in free range system and left to scavenge food whereas females are often better maintained, vaccinated, and reared in semi-intensive or intensive system due to litter needs.

Desi pigs reported a higher rate of infection than crossbreds because most of the desi breed pigs are reared in low input systems like free range or backyard farming, since they are comparatively hardy and easy to maintain and survive the temperature of state in better condition. Breeds like white Yorkshire are maintained in better health care systems like semi-intensive and/or intensive system and have better hygiene and low exposure to the infective eggs in the environment.

The current study observed least prevalence rate in intensive system of rearing 3.85% like research done in Punjab (19), supports the trends encountered in our study wherein only 0.50% (2/392) of prevalence rate was reported. A plausible explanation for the reduced prevalence in intensively managed pigs is the limited exposure to *T. solium* eggs, which are typically transmitted via fecal contamination. The intensive farming systems, restricts pigs to a controlled environments which involves confinement with regulated feeding, access to clean water sources, hygienic housing, and routine veterinary examinations. The above-mentioned conditions greatly reduce the risk of infection among pigs by encountering human feces — the primary source of *T. solium* eggs in endemic regions.

Moreover, biosecurity measures such as a comprehensive urban sanitation system involving a proper human waste disposal, disinfection protocols, and restricted access to free ranging pig helps to minimize environmental contamination.

These findings reiterate importance of clean environment and suggest’s transitioning from extensive to intensive farming practices could serve as an effective strategy in controlling porcine cysticercosis, thereby reducing the risk of transmission to humans and contributing to public health improvement, especially in endemic areas.

In conclusion, the observed lower prevalence in the intensive rearing system reiterates the importance of controlled and hygienic husbandry practices in breaking the life cycle of *T. solium* and underscores the need to promote such systems, particularly in high-risk regions.

A similar study in Peru (21), the product size observed by them varied considerably than what were observed in our study. In our study double bands were observed in microsatellite 27 and 32 (TSSR_27 & TSSR_32). For microsatellite 28 (TSSR_28) we observed size anywhere between 170-190 whereas in their study they observed a band width of 206-226. The polymorphism in band sizes across the two studies may be attributed to geographical and genetic variation among the *Taenia solium* isolates examined. Pajoulo and colleagues conducted their research in Peru, while the present study was carried out in India. Given the known genetic diversity of *T. solium* across different endemic regions, it is plausible that regional strains exhibit polymorphisms at specific microsatellite loci, resulting in variations in repeat lengths and, consequently, PCR product sizes.

Such genetic variability may reflect underlying evolutionary pressures, host-parasite interactions, and historical transmission dynamics that differ between continents. Microsatellite markers, being highly polymorphic and sensitive to mutation events such as insertions and deletions in repeat motifs, are particularly suited for revealing such intraspecies differences.

Therefore, the observed differences in microsatellite profiles between the two studies reinforce the importance of region-specific molecular epidemiological data in understanding the genetic structure of *T. solium* populations. Further, comparative studies need to be done using a broader set of markers and samples from diverse geographical regions would help elucidate the global genetic diversity of this parasite and enhance the accuracy of diagnostic and control strategies.

